# ProEnd: A Comprehensive Database for Identifying HbYX Motif-Containing Proteins Across the Tree of Life

**DOI:** 10.1101/2024.06.08.598080

**Authors:** David Salcedo-Tacuma, Giovanni Howells, Coleman Mchose, Aimer Gutierrez-Diaz, Jane Schupp, David M. Smith

**Affiliations:** Department of Biochemistry and Molecular Medicine, West Virginia University School of Medicine, 4 Medical Center Dr., Morgantown, WV USA; Department of Plant Biology, Uppsala BioCenter, Swedish University of Agricultural Sciences, Uppsala 75007, Sweden; Department of Neuroscience, Rockefeller Neuroscience Institute, West Virginia University, Morgantown, West Virginia, USA

## Abstract

The proteasome plays a crucial role in cellular homeostasis by degrading misfolded, damaged, or unnecessary proteins. Understanding the regulatory mechanisms of proteasome activity is vital, particularly the interaction with activators containing the hydrophobic-tyrosine-any amino acid (HbYX) motif. Here, we present ProEnd, a comprehensive database designed to identify and catalog HbYX motif-containing proteins across the tree of life. Using a simple bioinformatics pipeline, we analyzed approximately 73 million proteins from 22,000 reference proteomes in the UniProt/SwissProt database. Our findings reveal the widespread presence of HbYX motifs in diverse organisms, highlighting their evolutionary conservation and functional significance. Notably, we observed an interesting prevalence of these motifs in viral proteomes, suggesting strategic interactions with the host proteasome. As validation two novel HbYX proteins found in this database were tested and found to directly interact with the proteasome. ProEnd’s extensive dataset and user-friendly interface enable researchers to explore the potential proteasomal regulator landscape, generating new hypotheses to advance proteasome biology. This resource is set to facilitate the discovery of novel therapeutic targets, enhancing our approach to treating diseases such as neurodegenerative disorders and cancer. Link: http://proend.org/

## 1. Introduction

The proteasome is a key component in the cellular machinery, orchestrating the degradation of misfolded, damaged, or unnecessary proteins. Its function is critical not only for maintaining cellular homeostasis but also for regulating several biochemical pathways. In healthy cells, the proteasome ensures the proper turnover of proteins, preventing the accumulation of potentially toxic proteins that could disrupt cellular functions^1–3^. Conversely, in disease states, particularly in neurodegenerative diseases (ND) and certain cancers, the proteasome’s role becomes even more crucial. Aberrations in proteasome activity can lead to the accumulation of protein aggregates, a hallmark of many NDs, or affect the cell cycle and apoptosis, as observed in cancer^4–7^.

Because of this pivotal role/importance in disease, in the last decade several groups have been trying to generate drugs/pharmaceuticals to specifically target the proteasome and modulate its function in hope to present a promising strategy for therapeutic intervention ^7–9^. Velcade (Bortezomib), a pioneering proteasome inhibitor, exemplifies the proteasome’s potential as a drug target. Approved for the treatment of multiple myeloma and lymphoma, Velcade capitalizes on the need of proteasome activity for cell survival, particularly in cancer cells with high rates of protein synthesis and turnover^10–12^. This success increased the interest in researching and developing proteasome activators or alternative inhibitors that could modulate proteasome activity, offering new avenues for treating NDs. The activation of the proteasome, for instance, could enhance the clearance of protein aggregates, a shared feature of NDs, potentially improving disease symptoms or progression^7,8,13,14^.Thus, understanding the mechanisms of proteasome regulation, including the action of specific activators, inhibitors, and regulatory complexes, is vital for therapeutic intervention.

The 20S proteasome is defined as the core particle responsible for the proteolytic breakdown of proteins. Its cylindrical structure is composed of four stacked rings: two outer α-rings and two inner β-rings, each containing seven distinct subunits in the case of eukaryotes^15–18^. The catalytic activity resides within the β-rings, where three types of active sites, caspase-like, trypsin-like, and chymotrypsin-like facilitate the breakdown of proteins into peptides^15–17^. A critical aspect of the 20S proteasome’s function is its gate, formed by the N-termini of the α-subunits. This gate controls access to the inner chamber, where protein degradation occurs, hence regulating substrate entry. The gate’s default state is closed, preventing random protein degradation and ensuring selectivity in the proteasome’s degradative capacity^15–17,19^.

Another system for regulation of substrate entry is the existence of proteasome activators, such as the 19S regulatory particle (PA700), PA200, and PA28γ (also known as 11S), which bind to the 20S core and induce conformational changes that open the gate ^20–23^. The 19S regulatory particle, in particular, recognizes ubiquitinated proteins and uses ATPase activity to unfold and translocate them into the 20S core for degradation. Similarly, proteasome activators like PA200 or PA28γ enhance the proteasome’s ability to degrade specific non-ubiquitinated substrates, highlighting the versatility and adaptability of proteasome regulation^20,24^.

These mechanisms to regulate proteasome activity exhibit remarkable conservation throughout evolution, spanning from archaea to eukaryotes. This evolutionary continuity allows us to observe specific features that are essential across different domains of life ^25–27^. For instance, the archaea e.g. *Archegolbulous fuldgidous* contains a complete proteasome system including a 20S proteasome and an ATP-dependent proteasome activator known as PAN (proteasome-activating nucleotidase), which plays a pivotal role in this system, analogous to the function of the 19S regulatory particle in eukaryotic proteasomes^28^. Similarly, VCP (valosin-containing protein) also known as p97 or CDC48 in eukaryotes, has homologs in archaea (i.e. VAT) that shares functional similarities with PAN in its role in stimulating proteasome dependent protein degradation, highlighting the conserved nature of these proteolytic processes^29,30^.

The T20S proteasome (20S from *Thermoplasma acidophilum*) and its interaction with PAN have provided a simplified model to study the fundamental aspects of proteasome regulation. PAN, through its ATP binding activity, facilitates the opening of the 20S proteasome gate, enabling substrate entry, crucial for the degradation mechanism to function^31,32^. This interaction between PAN and the 20S emphasizes the importance of ATP-driven conformational changes for proteasome activity, a concept that is universally observed across species. Moreover, VCP/ p97 mediates the extraction of misfolded proteins from the ER for delivery to the proteasome for degradation, emphasizing the integrated nature of cellular degradation pathways^30,33–35^. Through the examination of the basal archaeal proteasome system and its eukaryotic counterparts, we have gained invaluable insights into the conserved mechanisms of proteasome degradation. Moreover, this evolutionary perspective increases our comprehension of cellular regulation but also highlights potential for discovery of therapeutic targets that span across species that await discovery.

The elucidation of the regulatory mechanisms governing the interaction between the proteasome activators and the 20S proteasome has been significantly advanced by the discovery of the HbYX motif. This tri-peptide motif, characterized by a hydrophobic amino acid (H), followed by a tyrosine (Y), and variable amino acids (X), which must be the C-terminal residue, has emerged as a key facilitator of proteasome activation and function. Pioneering work on the archaeal PAN complex and subsequent studies elucidated and highlighted the motif’s crucial role in gate opening, a necessary step for substrate entry into the 20S core^20^. The interaction of the HbYX motif with the 20S proteasome is mediated by its docking into specific pockets between adjacent α-subunits, known as α-pockets or intersubunit pockets, triggering allosteric changes that lead to gate opening, which have been recently characterized in mechanistic detail by Chuah et. al. 2023 and Gestwiki et al. 2022 36–38.

What makes the HbYX motif particularly intriguing is its prevalence among most proteins that bind to the proteasome^20^. This motif is not only conserved across a wide range of proteasomal ATPases, including those integral to the 19S regulatory particle and PAN, but also found in many proteasome regulators or proteasome binding proteins such as PA200, PI-31, Pac-1, Pac-2, and p97/CDC48/VAT. This conservation from archaea to humans highlight the motif’s fundamental role in proteasome binding and proteasome activation (gate-opening) denoting both, its utility and specificity in regulating protein degradation.

The question arises, can we identify other HbYX proteins to discover new pathways for protein degradation? In that context the use of bioinformatics has revolutionized our ability to uncover and characterize novel molecular entities, significantly advancing our understanding of complex biological systems. Protein domains and transcription factor motifs, crucial for regulating gene expression and protein function, serve as prime examples of targets identified through computational approaches^39,40^. However, the journey toward discovering specific proteasome regulators, and specifically those associated with the HbYX motif, presents unique challenges within the bioinformatics landscape.

Standard bioinformatics tools and pipelines, including the widely used BLAST or MEME (Multiple Em for Motif Elicitation) suite, offer powerful means to search for and analyze motifs within protein sequences^41,42^. These classical tools have been instrumental in identifying recurring patterns that play significant roles in protein interactions, localization, and function. Despite their utility, these tools often encounter limitations when tasked with detecting highly specific motifs like the HbYX, which only resides on the C-terminus of a protein. The inefficiency largely stems from the sheer diversity and complexity of protein length sequences, in addition, researchers often face challenges related to the structure of the data and the interpretation of outputs from these bioinformatics tools, requiring extensive manual curation to confirm the identified motifs.

In response to these challenges, there is a growing need for the development of specialized pipelines and tools^43^ tailored to the detection and analysis of specific proteasome regulators. Such advancements would enhance the efficiency and accuracy of motif discovery processes, enabling researchers to more effectively sift through the vast universe of proteins for potential therapeutic targets. Despite the hurdles, the potential to uncover novel regulators of the proteasome through bioinformatics and novel machine/deep learning methods remains a promising frontier in molecular biology and disease research.

In this report we introduce the **Pro**teasome **En**d regulators **d**atabase-ProEnd, a comprehensive database designed to navigate the universe of proteins for the discovery of new molecules featuring the HbYX motif. ProEnd, represents a validated tool to uncover and characterize the landscape of proteasomal regulators. ProEnd is specifically engineered to address the limitations encountered with standard motif search methodologies. By optimizing search to hone in on the unique characteristics of the C-terminal HbYX sequence, ProEnd has successfully identified a set of HbYX proteins across a wide range of organisms, from archaea to humans, constituting a specialized database dedicated to proteins with HbYX motif. This database serves as a foundational resource for researchers worldwide, facilitating searching, hypothesis generation, and exploration of the HbYX motif’s role in proteasome regulation and cellular function. By providing a platform for systematic analysis and cross-species comparison, we enable the scientific community to advance the understanding of new regulators and the diverse functions of the HbYX motif.

### Data collection and data processing

#### ProEnd in the UniProt/Swissprot reference universe

To address the question of the prevalence of HbYX motif-containing molecules and to overcome the limitations of existing bioinformatics tools, we developed a workflow designed to efficiently identify proteins with the specific C-terminal HbYX motif. Utilizing AWK, a text-processing tool, our pipeline operated through a three-step process that leverages protein data repositories for comprehensive analysis. First, we retrieved all reference proteomes from the UniProt^44^, representing all available spectrum of organisms. Subsequently, we streamline the proteome data into a simplified format, arranging it into a single file where each protein is listed on a separate line (Linearization). In the final step of our pipeline, we deploy a carefully designed regular expression to sift through the linearized array of proteomes. This search was finely tuned to detect the presence of the HbYX motif, which is only found on the C-termini of the protein with the C-terminal residue being the ”X” residue, ensuring that any occurrences are accurately captured. Upon identification, the protein alongside the motif sequence was cataloged in a structured table and we created a user-friendly online database (Fig1A) to allow researchers to interact with the information and structure prediction data. This approach not only simplifies the identification process but also facilitated a more manageable analysis of the data, allowing for a clearer understanding of the distribution and diversity of HbYX motif-containing proteins across different species.

### Technical validation

#### Analysis of HbYX proteins

Our comprehensive analysis covered approximately 73 million proteins across about 22,000 reference proteomes within the UniProt/Swissprot databases. The goal was to catalog all proteins featuring the HbYX motif. From this extensive survey, we observed an interesting and noteworthy phenomenon in viruses. While more than half of the viral proteomes examined do not possess any proteins with the HbYX motif, contrarily the other half of viral proteomes contain a high prevalence of HbYX motif-containing proteins compared to other organisms. Specifically, we identified approximately ∼7,500 HbYX proteins in viruses, which represent 1.92% of all proteins in these entities (Fig1B). Since a major function of the proteasome is to generate viral peptides for detection by the immune system (MHC-class I)^45,46^, these observations may indicate that virus could have evolved to either strategically avoid the proteasome by excluding HbYX motifs from their proteome or to negatively regulate it^47^.

In addition, bacteria, which do not have a proteasome system, presented approximately 235,000 HbYX motif-containing proteins, accounting for 0.72% of the proteins analyzed in these organisms. In bacteria, protein degradation is primarily performed by ClpP or HslV proteases and auxiliary AAA-ATPase particles such as ClpA, ClpX, and HslU. These complexes are compartmentalized proteases like the eukaryotic proteasome but are not evolutionarily related^48,49^. Although some studies do suggest HslU/V is somewhat related to the proteasome system, and the C-terminus of HslU does bind to HslV in an ATP-dependent manner for regulatory purposes, which is analogous to 19S-20S or PAN-20S association^50,51^. However, HslU does not have a HbYX motif ^50,51^. In addition, as a group, these bacterial protease systems typically rely on adaptor molecules to target proteins for degradation^52,53^. The presence of HbYX motifs within bacterial proteins suggests a broad, cross-kingdom utility for these sequences in protein interaction networks. A notable example is found in Actinobacteria, which have acquired a eukaryotic-like 20S proteasome through horizontal gene transfer. This proteasome is regulated by a prokaryotic ubiquitin-like protein (Pup) and an ATPase cap (Mpa) that contains a variant of the HbYX motif and is structurally similar to eukaryotic AAA-ATPases^54,55^. In Actinobacteria, we identified 982 proteins that feature the HbYX motif (Fig1B).

Archaeal proteomes revealed ∼7.2 thousand proteins with the HbYX motif at their C-termini, corresponding to 0.93% of the proteins in these foundational life forms. This finding highlights the evolutionary continuity, as archaea, like eukaryotes, rely on the proteasome system for protein degradation. The archaeal proteasome system consists of a simpler 20S catalytic core and a hexameric unfoldase complex known as PAN. Similar to the eukaryotic 19S ATPase complex (Rpt1-6), the PAN AAA-ATPase employs mechanochemical forces to unfold and translocate target polypeptides into the 20S catalytic chamber^26–28^. Like all known 20S caps in eukaryotes and actinobacteria, PAN features a conserved C-terminal hydrophobic-tyrosine-X (HbYX) motif that facilitates proteasome gate opening and substrate entry^26–28^. We identified 60 different PAN proteins across the archaeal proteomes (Fig-1C). Furthermore, the VCP/p97/VAT/CDC48 AAA-ATPase, which also contains the HbYX motif, has been shown to interact with the archaeal 20S proteasome and may function as an alternative regulatory complex, although its binding to the eukaryotic 20S proteasome and its effectiveness as a proteasome cap are still under debate^30,33,35^. We identified 54 VAT/CDC48 proteins with the HbYX in archaeal proteomes. For perspective, the single archaea organism, *Archaeoglobus fulgidus*, has 44 HbYX proteins including one CDC48, but most of which have unknown functions. The detection of HbYX motif-containing proteins in archaea suggests a possible regulatory network conservation of activators related to this motif across these domains.

Eukaryotic organisms displayed approximately 405 thousand proteins with the HbYX motif, about 1.01% of their total protein count (Fig1B). This is expected given the complexity and specialization of the eukaryotic proteasome system, which needs diverse interactions with various degradation pathways. From this amount 3,266 are known activators from the 19S family of regulators and PA200 (Fig-1C). In humans specifically, we found 234 proteins with the HbYX motif, representing 1.67% of the protein-coding sequences. This is notably lower than the anticipated random frequency of 2.5% for any given amino acid sequence, implying a potential evolutionary pressure to constrain the abundance of HbYX motif proteins, possibly due to their specialized function in proteasome regulation. Prior studies have also attempted to identify Proteasome Interacting Proteins (PiPs) by various methods such as pulldowns and mass spec. When we compared the known PiPs with all identified human HbYX proteins in ProEnd we found that 22.2% of these human HbYX proteins have indeed been identified previously as PIPs ^56–58(Fig1D)^. These findings provide independent validation to the idea that HbYX proteins have a high probability to bind to the proteasome. This validation is extended to the archeal organism *Thermoplasma acidophilum,* a widely known model organism used to understand proteasome function. In this organism independent validations have shown that 25% of the HbYX containing proteins have been detected by pulldown with the 20S proteasome^30^ (Fig1D). These independent validations show the potential of the HbYX motif as a cross-kingdom characteristic that indicates the ability to interact with the proteasome and potentializes all proteins containing HbYX in the C-terminus as PiPs, many of which may be modulators of proteasome function or related degradation pathways.

### Biological-driven hypotheses generation for HbYX-containing proteins

The most well-known and studied function of the HbYX motifs is its ability to bind to and activate the 20S proteasome. Known regulators such as PA200, PI31, and PACs, are capable of activating, inhibiting, and assembling the proteasome, respectively^22,59,60^. To determine additional functional diversity associated with the HbYX motif identified by our approach we performed a gene enrichment analysis^61^ in model gnathostomata organisms. Among jaw vertebrates the enrichment revealed a prevalence of HbYX proteins in transmembrane transport, neurotransmitter activity, and GABA receptor-related functions (Fig 2A). In humans, from the pool of HbYX proteins, seven have been identified as involved in Alzheimer’s disease pathways based on literature (not counting the proteasome itself), and all but one bind APP and include Presenillin 1 & 2^62^. Interestingly, Presenilin’s HbYX motif is extracellular, interacts with a pocket on the gamma-secretase subunit^63^ (Fig 2B), affecting its substrate gating and it is highly conserved among in jaw vertebrates (Fig 2C). This suggests that HbYX motifs may regulate transmembrane enzymatic function/activity and could potentially recruit extracellular proteasomes when not bound to transmembrane partners. The 20S is present in the cerebrospinal fluid, indicating putative extracellular 20S docking sites with unknown functions. Most interestingly we found that GPM6A, which has been suggested to associate with Neuronal Membrane Proteasomes (NMPs^64^) is a HbYX protein and its motif (AYT) is conserved among jawed vertebrates (Fig 2D-E). The NMP is emerging as an important factor in regulating neuronal activity and is proposed to be associated with neurodegeneration^64,65^. The existence on NMP open the possibilities for the HbYX proteins found in the neuronal context such as GABA receptors and others in our database as potential recruiters or modulators of proteasome function in different cellular contexts from those expected so far, where protein degradation by the proteasome remains uncharted territory. Further validations are necessary to fully understand the context and function of these molecules and whether there is a proteasome-independent function for HbYX molecules, which seems likely, or if is recruited to perform localized functions in cellular context. These are just a few examples of the potential hypotheses generated from our HbYX protein database, as HbYX-proteins can be found in almost every cellular compartment, such as cytosol, nucleus, ER, Golgi, nucleoplasm, mitochondria, peroxisomes, and nuclear speckles.

**Figure 1.**
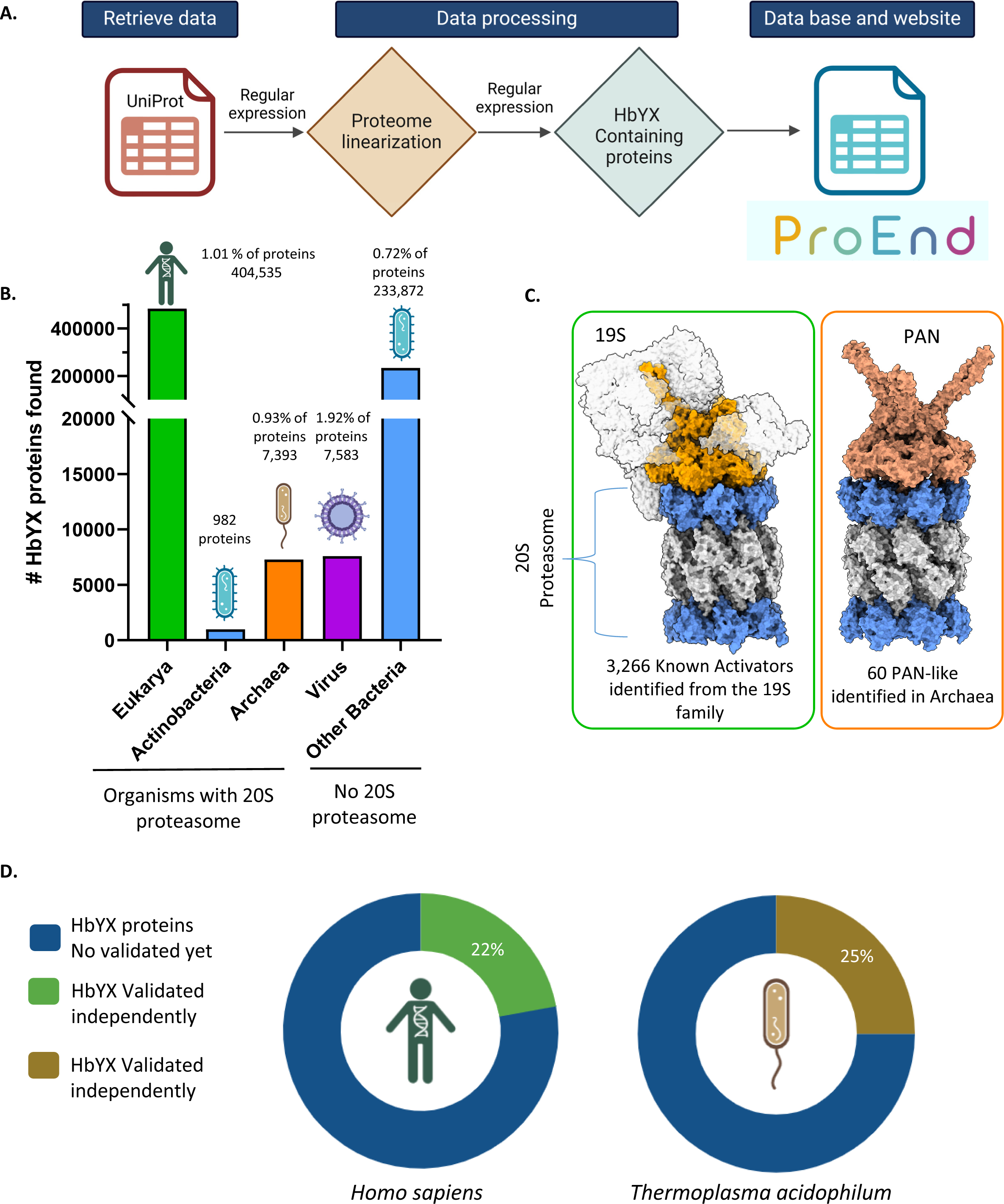
Prevalence of HbYX motif across organisms identified by ProEnd. **A.** Workflow for the design of the ProEnd database and the pipeline used to retrieve proteomes from UniProt/SwissProt. AWK was utilized to develop the regular expressions. **B.** Distribution of HbYX Proteins Across Life Domains and Viruses. Barplot indicates the number of HbYX proteins within various life domains, represented as percentages indicating the average occurrence of HbYX proteins per domain. On the left side, organisms possessing a 20S proteasome—specifically Eukaryotes, Actinobacteria, and Archaea—are grouped as highlighted by the line below. On the right, organisms lacking a 20S proteasome, including viruses and other bacteria, are displayed. **C.** Number of known proteasome activators found in the ProEnd database. Structural representation of a 20S proteasome with α-rings in blue and β-rings in gray. The green box illustrates the eukaryotic 19S regulatory particle complexed with the 20S proteasome (PDB: 6MSB), highlighting the conserved ATPases in orange. Additional regulatory proteins are outlined to enable comparison with the archaeal proteasome regulator. The orange box displays the structural complex of the archaeal 20S proteasome and PAN (PDB: 6HEC), with the PAN cap shown in salmon. In total 3,266 known activators form the 19S were identified in Eukarya and 60 PAN-like proteins were identified in Archaea. **D.** Percentage of HbYX proteins independently validated as proteasome interactors in other studies for *Homo sapiens* and *Thermoplasma acidophilum,* an archaeal model organism.

**Figure 2.**
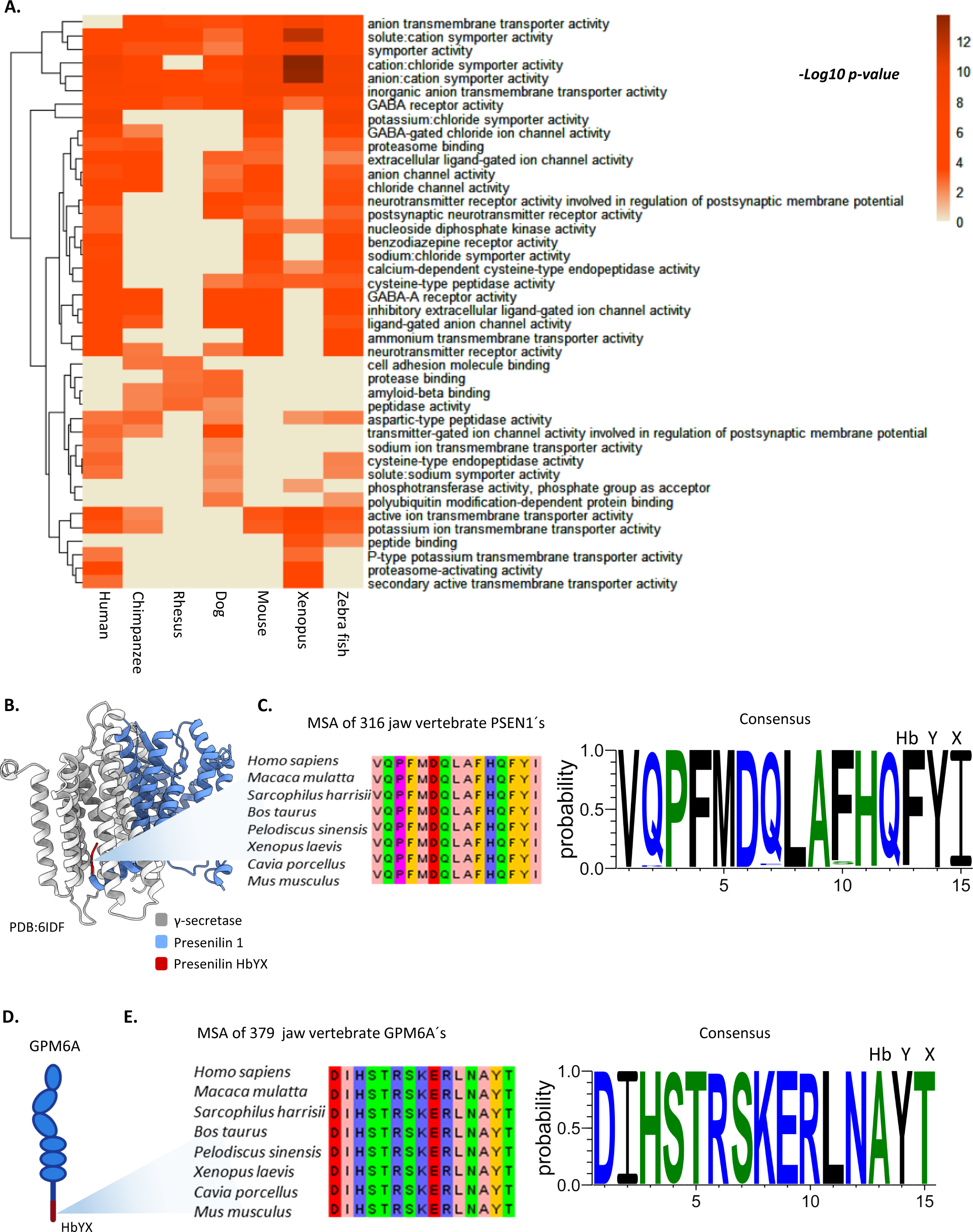
Conserved enrichment of HbYX proteins associated with neuronal pathways among jawed vertebrates indicates conserved functions for the HbYX motif. **A.** Heatmap representing molecular function enrichment among model jawed vertebrates. The heatmap is colored by the significance of enrichment (’-log10 p-value’). Higher values indicate increased significance associated with the pathway. Rows indicate the enriched processes, and columns represent the model animals. Enrichment was determined using ClusterProfiler. Zero values indicate no enrichment found for this process among the HbYX proteins. **B.** Presenilin interaction with γ-secretase (PDB: 6IDF). This model represents the HbYX interaction of PSEN1 with γ-secretase, suggesting a potential binding role of the HbYX motif. γ-secretase is shown in gray, PSEN1 in blue, and the HbYX motif in red. **C.** Conservation of the HbYX motif in jawed vertebrates. A total of 316 sequences of PSEN1 were recovered from different vertebrates, and a multiple sequence alignment (MSA) was constructed using Jalview. On the right, the hidden Markov model (HMM) logo for the consensus sequence of the 316 PSEN1 sequences shows the high conservation of the HbYX motif (FYI). **D.** Representation of the neuronal membrane glycoprotein GPM6A and its HbYX motif. **E.** Conservation of the HbYX motif in jawed vertebrates A total of 379 GPM6A sequences from jawed vertebrates were retrieved, and an MSA was constructed using Jalview, highlighting the conserved features of this protein. The consensus HMM logo was then plotted, showing a highly conserved HbYX motif (AYT).

### Utilizing ProEnd for investigating Proteasome Interactions with HbYX Proteins

To validate the utility and usage of ProEnd for novel proteasome interacting proteins, we tested 2 examples from different organisms. The first one we selected related to neuronal regulation is an interesting example of a highly conserved protein among jawed vertebrates, neurotensin, a small protein that contains the HbYX motif in the form of YYY. This motif is conserved across all vertebrates with minimal changes in *sauropsides* where the initial Y is replaced with a S (Fig 3A). Under physiological conditions Neurotensin is a regulatory peptide of 13 amino acids originated by the cleavage of the full length precursor protein, predominantly located in the gut and brain influencing various dopaminergic pathways^66,67^. The cleavage of the neurotensin precursor results in the release of neurotensin and neuromedin N, along with other peptide fragments. These peptides are less potent than neurotensin but are also less sensitive to degradation, which may allow them to act as longer-lasting modulators in various physiological situations^68,69^. One of these fragments is the very C-terminal which contains the HbYX YYY motif. To test whether it is associated with the proteasome, human biotin-fused to neurotensin C-terminal HbYX peptide was synthesized to perform a pull down assays on HEK293 cell lysates. Our results showed that neurotensin can pull down the 20S proteasome from human cells (Fig 3B-C), indicating an interaction between this highly conserved HbYX-protein and proteasome degradation pathways. This suggests that highly conserved proteins with the HbYX motif are capable of binding to the proteasome, hence, its function, conserved through evolution can be related to proteasome binding and regulation. It’s important to note that, to date, there is no evidence in the literature suggesting that the neurotensin peptide or other HbYX motif-containing proteins related to neurotransmission, such as GABA receptors, are associated with proteasome binding, revealing that many of the HbYX molecules identified could recruit the proteasome to execute functions in different cellular contexts, increasing the number of functions associated with the proteasome and opening the door to new possibilities for understanding the cellular context of degradation in each cell type. Neurotensin presents an example of how proteins found with HbYX motif in ProEnd database uncover a rich niche of new functions, hypotheses, and approaches to understanding proteasome biology.

**Figure 3.**
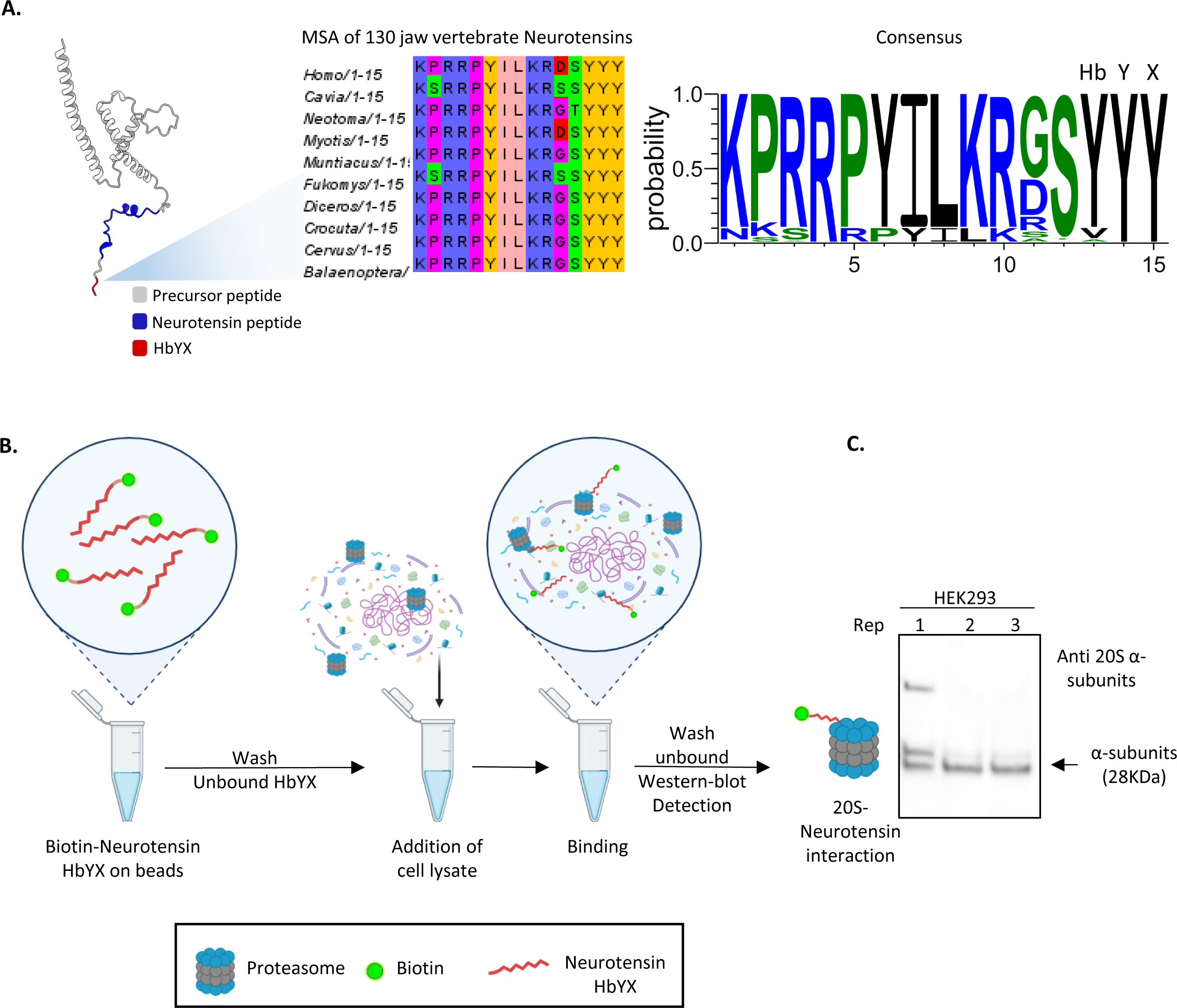
Neurotensin as a conserved HbYX-containing protein with the ability to bind the proteasome. **A.** AlphaFold prediction of the neurotensin precursor (AAF-P30990-F1) showing the Neurotensin peptide in blue and the HbYX motif in red. For the HbYX motif, a multiple sequence alignment (MSA) was constructed using neurotensin sequences from 130 vertebrates to show conservation of the HbYX motif. The consensus for the HbYX motif was constructed using an HMM logo, revealing a highly conserved YYY motif, with occasional occurrences of valine and alanine in the hydrophobic position. **B.** Validation pull-down assay performed with the Neurotensin HbYX peptide fused to biotin. Briefly, biotinylated Neurotensin was incubated with avidin beads. After washing off unbound HbYX, cell lysates from HEK293 cells were added for binding. Following incubation, unbound proteins were washed off, and detection of the proteasome was performed through western blotting for the alpha subunits. **C.** Western blot for the pull-down assay described in B, probing for alpha subunits with an expected molecular weight of 28 kDa. Three different replicates of HEK293 cell pull-downs were tested, showing the presence of alpha subunit proteasome bands. This indicates that the neurotensin HbYX peptide can pull down proteasomes, demonstrating that neurotensin is a proteasome interactor protein. The upper band in replicate 1 indicates the presence of the 26S proteasome.

Our second example comes from archaea, highlighting the proteasome activities mediated by HbYX proteins in these organisms. Specifically, we investigated a protein from *Methanocaldococcus jannaschii*, this specie has the known proteasome activator PAN^70^, but with ProEnd help we identified a HbYX containing protein annotated as putative 26S proteasome regulatory subunit by similarity according to UniProt (MJ1494). Motivated by its potential role as a proteasome regulator, we generated this HbYX protein recombinantly to assess its regulatory capabilities within the proteasome. To assess its ability to bind and regulate the archaeal proteasome T20S, we employed an experimental approach similar to Fig 3B but with slight modifications. First, recombinant T20S was purified using Ni-NTA affinity chromatography followed by TEV cleavage in order to remove the 6His tag. Concurrently, recombinant HbYX protein MJ1494 was purified through sonication and Ni-NTA affinity chromatography. Prior to elution, purified and TEV cut T20S was added to columns containing HbYX-bound MJ1494 and washed thoroughly. The HbYX protein was then eluted using wash buffer supplemented with 150mM imidazole. The presence of T20S in the elution fractions was determined by a proteasome activity assay using the fluorogenic substrate suc-LLVY-AMC, which is cleaved by the T20S. The specificity of this assay allows us to measure the presence of T20S accurately and to calculate the initial velocity (Vo) of substrate degradation, unaffected by any regulatory effects on the T20S gate, since gate-closure does not affect LLVY degradation. We found that MJ1494 pulled down 7X more T20S, than did control Ni-NTA columns (Fig 4A). This result validates MJ1494 as a HbYX containing protein with the ability to bind to the proteasome, adding another validation to support the HbYX containing proteins found in our ProEnd database as proteasome regulators. Additionally, to understand MJ1494 structure conformation we predicted its homo-oligomerization using AlphaFold Multimer^71^. We expected homo-oligomerization since MJ1494 contains an ATP binding domain in the 161-168 positions, and it is expected, based on homology with ATPases such as PAN, to form hexamers. The AlphaFold prediction suggested a hexameric assembly with a pLDDT score of 0.78 (Fig 4B, 4D HbYX in orange), indicating good overall structural accuracy. The Predicted Aligned Error (PAE) plot revealed high confidence in the intra-monomeric contacts and moderate confidence in the inter-monomeric interactions, consistent with the hexameric structure (Fig 4C). While the local structures are reliable, some inter-chain interactions may require further experimental validation. According to the AlphaFold prediction the HbYX motif seems to be buried in the ATPase ring which is also a common feature shared with PAN when it is not bound to ATP and not bound to the proteasome, indicating structural similarities between PAN and MJ1494. Noteworthy is that MJ1494’s homology percentage to *M.jannaschii*, PAN is 29.77% using the BLOSUM62 matrix. While this percentage suggests some sequence similarity, it is relatively low, indicating a distant evolutionary relationship and potentially different regulatory roles or specificities. Importantly, MJ1494 does possess the N-terminal OB domain, akin to other proteasome ATPases, suggesting a conservation of structural features despite the low homology. This finding enhances our knowledge of proteasome activators in ancient organisms like archaea, demonstrating the variability in PAN-like proteins across distinct organism families and it also illustrates the evolutionary adaptation and functional differentiation of proteasome regulators in diverse biological contexts.

**Figure 4.**
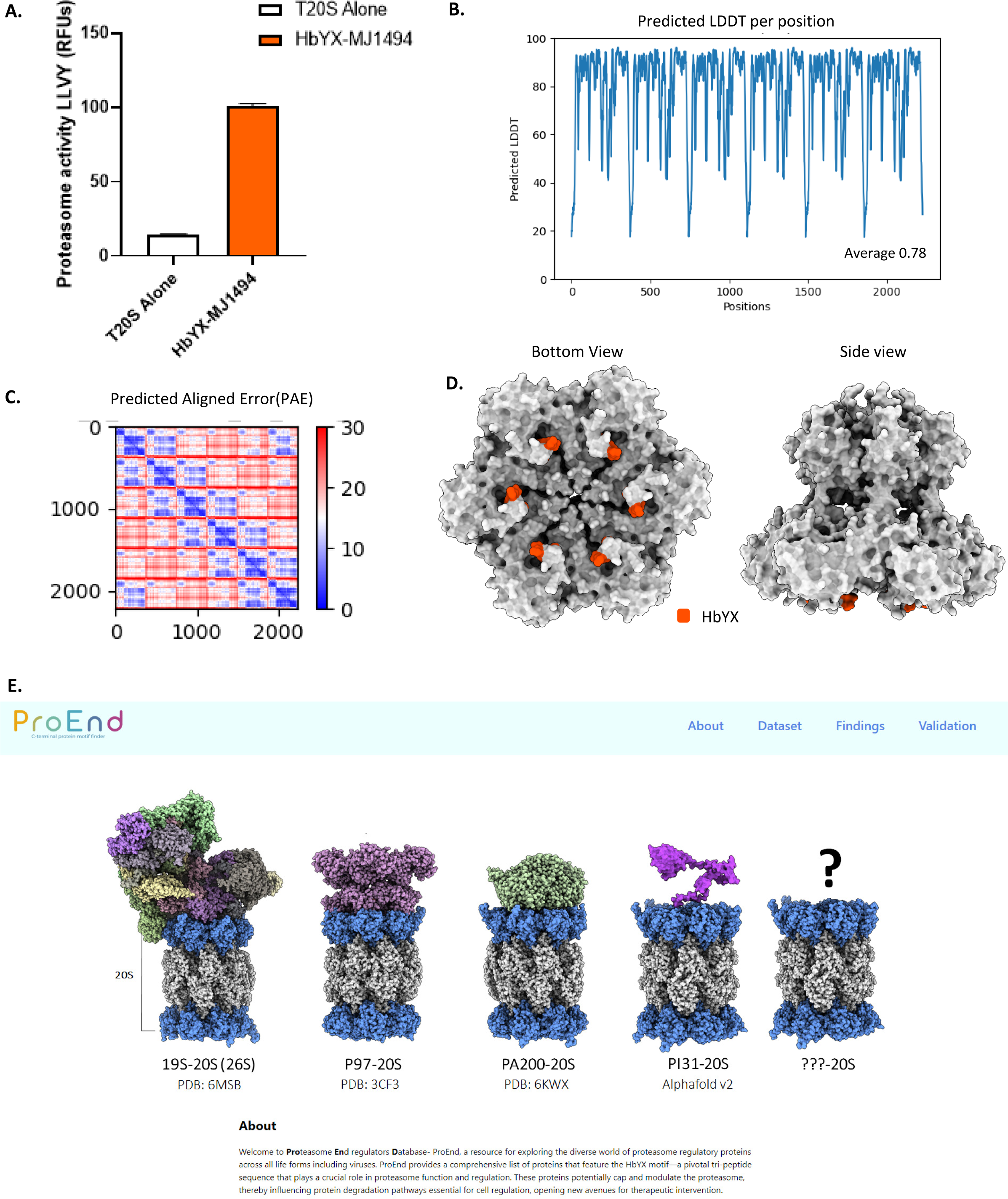
Archaeal HbYX ATPase MJ1494 is a PAN-like protein that binds and activates the archaeal proteasome T20S. **A.** Rate of substrate degradation of fluorogenic LLVY after pulldown with T20S on MJ1494 , pulldown with proteasome activator PA26, or T20S alone. Stimulation of degradation was measured by the increase in LLVY hydrolysis (rfu). **B.** AlphaFold multimer prediction LDDT indicating overall confidence in the predicted structure. Scores above 0.51 are generally accepted. Scores above 0.70 are considered to be of good quality, suggesting that most regions of the protein are predicted with high accuracy. **C.** Predicted Aligned Error (PAE) provides insights into the accuracy of the predicted inter-residue distances. The diagonal blocks with lower PAE values (indicated by blue regions) suggest that the intra-monomeric contacts are predicted with high confidence. This aligns with the high pLDDT score (0.78), indicating reliable local structure predictions within each monomer. The off-diagonal blocks show higher PAE values (indicated by red regions), reflecting moderate confidence in the inter-monomeric contacts, indicating some uncertainty in the predicted interactions between monomers. The repeating pattern of the blocks in the PAE plot is consistent with the hexameric nature of the protein, suggesting that each monomer’s predicted structure is similar and forms repetitive interactions within the hexamer. **D.** Bottom and side views of the structural representation predicted by AlphaFold multimer visualized with ChimeraX. The complex is shown in gray with the HbYX motif highlighted in orange for visualization. The HbYX motif appears to be buried inside the ATPase ring, a common feature shared with PAN when not engaged with the 20S proteasome. **E.** Screenshot of the main page of ProEnd. The database is available at http://proend.org/

### A web portal for HbYX motif-containing proteins

To allow researchers to interact with the comprehensive dataset of HbYX motif-containing proteins, we created ProEnd, a user-friendly online database (first released in 2024 Fig 4E). Users can explore an expandable, interactive matrix or a sortable list that can be filtered by species, domain, or gene name. Clicking on a matrix tile or list entry displays an information page that includes an interactive protein structure viewer, where AlphaFold structure predictions of the HbYX motifs can be examined and downloaded. The information page also provides UniProt entry information. These features facilitate rapid searching, visualization, and analysis of several HbYX proteins, making ProEnd an invaluable resource for the global scientific community. The database is hosted on Amazon Web Services using MongoDB, ensuring high availability and performance. Researchers can access ProEnd at http://proend.org/ and navigate the data to support their studies in proteasome regulation and beyond.

## Conclusions

The establishment of the ProEnd database marks a significant advancement in the field of proteasome biology, providing an invaluable resource for identifying and characterizing HbYX motif-containing proteins across a broad spectrum of species. By offering a comprehensive catalog and an interactive platform, ProEnd facilitates the exploration of potential proteasomal regulators and their evolutionary conservation, uncovering their potential diverse roles in cellular processes. The findings reported here highlight the prevalence and importance of HbYX motifs in proteasome regulation, with implications extending from basic biological research to potential therapeutic applications. This database is expected to not only enhance our understanding of proteasome mechanisms but also opens new avenues for the discovery of novel proteasome modulators, with potential far-reaching impacts to understanding proteasome biology and treating human disease.

## Methods

### Enrichment analysis

Enrichment analysis was performed for jawed vertebrate species by extracting lists of HbYX proteins and analyzing them in R (version 4.1.2) with the ’clusterProfiler’ package^61^. The ’enrichGO’ function was employed for each species, and a heatmap was created using - log10 transformed data to highlight significant enrichments in molecular functions.

### MSA and HMM logo construction

Sequences of HbYX proteins were identified with ProEnd, retrieved from the UniProt database and aligned using Jalview^72^ with the ClustalX alignment tool. For the construction of Hidden Markov Model (HMM) profiles, we utilized WebLogo3^73^ and Python library LogoMaker to generate sequence logos and visualize the conservation and variability of the HbYX amino acids. For MJ1494, the substitution matrix BLOSUM62 was calculated using Python Bio.Align to assess evolutionary divergences and similarities with PAN.

### HbYX containing protein plasmid construction and Proteasome purifications

The DNA sequence for the HbYX protein MJ1494 was fetched from UniProt and cloned into pET-28a(+) vectors for recombinant *E. coli* overexpression under a T7 promotor and lac operator. A 6His tag was added to the N-terminus of each construct for Ni-NTA purification and pulldown experiments. The 6His tag is followed by a TEV protease sequence (ENLYFQ/M) where “/” indicates the cut site. The pET28a(+) vector was chosen due to stable expression of other archaeal proteins in the past in our lab. DNA sequences were codon optimized for expression in the bacterial strain BL21 star DE3. Transformed bacteria are overexpressed through IPTG induction. *T. acidophilum* 20S proteasomes were expressed using the pET-28a(+) vector and contain a C-terminal 6His tag on the β subunits preceded by a TEV protease sequence in order to remove the 6His tag following purification. The T20S was purified following Seemüller et. al ^75^, with variations. Following purification, the T20S was TEV cut. Following TEV cleavage the T20S was added to an Ni-NTA column and only the flow through collected ensuring all T20S used in pulldown experiments are tagless.

### Proteasome activity assays–peptide substrates

Fluorogenic substrate peptide, suc-LLVY-amc, was obtained from BostonBiochem. For activity assays, this substrate was dissolved in DMSO and used at a final concentration of 100 µM. The reaction mixture also included a final DMSO concentration of 2%. The assays were performed in a buffer at 45°C, with 2 mM ATP and 10 mM MgCl2, to a total volume of 0.1 ml of reaction buffer. The assays were conducted over 30 to 60 minutes and the results were analyzed using BioTek Gen5 Data Analysis software. Fluorescence at excitation/emission 380/460 nm was measured every 55 seconds for 2 hours. As LLVY bypasses the T20S gate the initial rate of increase in fluorescence intensity is directly proportional to proteasome quantity and activity.

### Cell culture

HEK-293 cells were grown in DMEM (Life Technologies, Carlsbad, CA) supplemented with 10% FBS (Sigma, St. Louis, MO), and 1% penicillin/streptomycin (Life Technologies). Cells were cultured at 37°C in 5% CO2 environment at a density of 5x106 cells/ml and collected to for pulldown assays. All cells were obtained from ATCC (Manassas, VA).

### Ni-NTA pulldown assay

Cell lysates were clarified by centrifugation at 20,000rpm using a F21-8x50y rotor (ThermoFisher) for 30 minutes. Next, 200µL of Ni-NTA agarose resin (Qiagen) is added to the supernatant and allowed to bind for 1 hour shaking at room temperature. The resin is washed extensively with wash buffer (50mM Tris + 100mM NaCl supplemented with 2mM ATP, 10mM MgCl2. Following this washing step the columns are capped and concentration matched T20S is added and allowed to incubate for 20 minutes at room temperature. After 20 minutes the column is uncapped and the flow-through of unbound proteins is allowed to flow through the column. Columns are then washed with buffer (1x column bed volume) to further remove unbound proteins. Finally, HbYX containing proteins along with HbYX bound proteins are eluted using the wash buffer supplemented with 150mM imidazole. Presence of 20S in the elution fractions indicates binding to the HbYX protein bound on the Ni-NTA column, and the amount of 20S is quantified in the elution fraction by running a proteasome activity assay using the fluorogenic substrate suc-LLVY-amc as well as SDS-PAGE and western blot in the case of neurotensin.

### SDS-PAGE and Western blot

Proteins were separated by SDS-PAGE using Invitrogen NuPAGE 4-12% Bis-Tris protein gels following manufacturer’s protocol. Samples were mixed with Invitrogen SDS sample buffer (4x). Protein bands were visualized with Coomassie stain (Simply Blue Safe Stain, Novex) according to manufacturer instructions. Protein band sizes (kDa) was visualized using the SeeBlue Plus2 pre-stained protein standard (ThermoFisher). For western blot, the gel was transferred to an Immobilon®-FL PVDF membrane at 30V overnight. The membrane was blocked in Tris-buffered saline with 0.01% Tween-20 (TBST) and 10% nonfat milk for 1 hour at room temperature (∼20°C), then briefly washed and incubated with the primary antibody (Anti-Proteasome α’s1,2,3,5,6,7 subunits, Enzo, MCP231; Anti-PSMD7/Mov34, ab140428) diluted 1:1000 in TBST with 5% nonfat milk at 4°C overnight. After washing three times for 5 minutes each with TBST, the membrane was incubated with the secondary antibody (DyLightTM 550, Thermo, 10173; Alexa Fluor Plus 680, A32729) diluted 1:3000 in TBST with 5% nonfat milk for 1 hour at room temperature. The membrane was washed again three times for 5 minutes each with TBST and then imaged using an Amersham Typhoon (GE).

### Alphafold Multimer predictions

Structural models of MJ1494 (UniProt ID: Q58889) were generated using the AlphaFold Multimer ^71^ v2.3.295, specifically configured to predict multimeric assemblies, reflecting the protein’s anticipated homohexamer formation. This computational modeling was performed on a high-performance computing platform, utilizing six chains to represent MJ1494’s oligomeric state accurately. The analysis employed a reduced database preset, with a template search cutoff incorporating the most current structural data available up to August 1, 2023, ensuring the model’s relevance with the latest updates. The resulting models were visualized and refined using ChimeraX software.

## Acknowledgements

We extend our appreciation to the members of the Smith Lab for their invaluable discussions and feedback, which significantly enriched the development of this manuscript, particularly to Dr. Chuah for her expert insights. We also owe a special thanks to Dr. Chris Adami and the team at the Institute for Cyber-Enabled Research at Michigan State University. Their computational resources and services were instrumental in enabling the AlphaFold oligomer modelling. Lastly, we are grateful for the financial support from the NIH, specifically grants R01AG064188 and R01GM107129 awarded to D.M.S., which made this work possible.

